# Dysregulated Platelet GPIbα–VWF Signalling in Abdominal Aortic Aneurysm formation and Progression

**DOI:** 10.64898/2026.09.21.751373

**Authors:** T Feige, KJ Krott, A Bosbach, M Chario, J Ortscheid, A Saleem, MU Wagenhäuser, H Schelzig, M Elvers

**Affiliations:** Department of Vascular- and Endovascular Surgery, University Hospital Duesseldorf, Heinrich-Heine-University, Duesseldorf, Germany

**Keywords:** platelets, glycoprotein Ib, abdominal aortic aneurysm, ECM remodelling

## Abstract

**Background:** Platelets are critical drivers of thrombo-inflammatory responses in different cardiovascular diseases. Abdominal aortic aneurysm (AAA) is a progressive, life-threatening vascular disorder mainly characterised by chronic inflammation, extracellular matrix degradation, and the formation of a platelet-rich intraluminal thrombus (ILT). Experimental and clinical evidence identified platelets as main players in AAA pathology as evidenced by elevated platelet activation and procoagulant activity that critically contribute to AAA progression.

**Methods:** The present study investigated the contribution of glycoprotein (GP)Ibα, the von Willebrand factor (VWF)-binding subunit of the platelet GPIb–IX–V complex, to AAA initiation and progression in experimental AAA using the ePPE mouse model and in patients.

**Results:** Genetic ablation of platelet GPIbα significantly attenuated early aneurysm expansion in experimental AAA, indicating a critical role for GPIbα during the initial stages of aneurysm development. This initial effect was compensated at later time points showing no differences in aneurysm progression between groups. Notably, genetic deletion of GPIbα induced a constitutively hyperactive platelet phenotype already in naive mice that was further amplified during experimental AAA. This elevated platelet hyperactivity was mainly due to increased GPVI activation of platelets 28 days post-surgery. To assess the clinical relevance, spatial profiles of human ILT specimens from patients with AAA were analysed. In the ILT, we detected a highly compartmentalised distribution of GPIbα and VWF with pronounced enrichment within the luminal layer. In parallel, circulating VWF activity as well as platelet surface expression of GPIbα were significantly increased in patients with AAA.

**Conclusion:** Collectively, these findings identify a dysregulated GPIbα–VWF axis in human AAA pathology, mainly characterised by enhanced platelet GPIbα surface expression and increased activity of circulating VWF.

## 1. Introduction

Abdominal aortic aneurysm (AAA) is defined by a pathological and progressive dilatation of the abdominal aorta [1]. Established risk factors include advanced age, male sex, hypertension, smoking, a positive family history of AAA, elevated cholesterol levels, and pre-existing cardiovascular comorbidities [2]. On a global scale, AAA is ranked among the top ten causes of cardiovascular-related deaths [3]. Given the pronounced age-dependent increase in prevalence, AAA affects approximately 0.92% of individuals aged 30 to 79, corresponding to an estimated global burden of 35 million diagnosed cases in 2019 [4]. From a pathophysiological perspective, AAA formation is characterised by chronic inflammation, extensive vascular remodelling, and the formation of a platelet- and RBC-rich intraluminal thrombus (ILT) [1, 5–7]. In this context, AAA progression is mainly driven by enhanced platelet activation, increased procoagulant activity, and infiltration of inflammatory cells into the aortic wall. These mechanisms in turn facilitate expression and activation of proteolytic enzymes – such as MMPs – reinforcing the progressive degradation of extracellular matrix components. This culminates into a compromised aortic wall integrity within the aneurysm segment, consequently leading to an irreversible dilation of the affected aortic tissue [8]. Thereby, this continuous expansion is closely associated with fatal aortic rupture, which substantially contributes to the high mortality rate observed within AAA pathology. Despite considerable efforts during the last decades, the underlying molecular and cellular mechanisms of AAA pathogenesis are still under the current scope of investigation, especially as no pharmacological therapeutic intervention to effectively restrict aneurysm growth is available to date. Although multiple pharmacological approaches have demonstrated promising results in preclinical settings, their translation into a clinical application is still lacking and remains challenging, as several clinical trials failed to yield convincing evidence of clinical efficacy [9–12].

Besides their fundamental role in haemostasis, platelets act as important drivers of pathological processes in different cardiovascular disorders, including myocardial infarction, stroke, and AAA [13, 14]. In this context, cumulating experimental evidence implicates platelet-driven mechanisms as critical contributors to the development and progression of AAA. In line with this, clinical data highlight that application of low-dose aspirin may offer protective effects in patients with AAAs exceeding 35 mm in diameter, thus supporting the potential of antiplatelet therapies as pharmacological intervention, especially in advanced disease stages [15]. Notably, these beneficial effects have not been observed in small-sized aneurysms. However, these contradictory results clearly underscore the urgent need for a more comprehensive understanding of the underlying platelet-dependent mechanisms, in order to identify novel and promising platelet-specific targets [16–22]. In this context, Wagenhäuser *et al*. recently identified platelets as key drivers of aneurysm formation by modulating both inflammatory responses and structural remodelling of the aortic vessel wall. Mechanistically, platelet depletion in an *in vivo* mouse model of experimentally induced AAA formation, resulted in a pronounced suppression of key inflammatory genes – such as *Il1b*, *Il6*, *Il8*, *Il10*, *Il12*, and *Tnf-*_α_ – and extracellular matrix remodelling genes – including *Mmp9* and *Col1a1* – within the aneurysmal tissue, thereby significantly attenuating aortic diameter expansion. In addition, comprehensive phenotypic analyses of platelets derived from patients diagnosed with AAA revealed a persistent hyperactive state under basal conditions, as reflected by increased degranulation (P-selectin surface expression) and enhanced integrin α_IIb_β_3_ activation. This was paralleled by substantially elevated procoagulant activity, indicated by increased phosphatidylserine (PS) exposure, thus pointing to a platelet-driven prothrombotic phenotype in AAA [8, 19]. The main platelet collagen receptor glycoprotein (GP) VI has been identified to trigger platelet hyperactivity in AAA and thus emerged as a platelet-specific target for a promising therapeutic approach [16, 23]. In detail, genetic deletion of GPVI substantially reduced experimental AAA *in vivo*, primarily by limiting neutrophil recruitment and subsequent neutrophil extracellular trap (NET) formation, while concurrently attenuating extracellular matrix remodelling via downregulation of MMPs. These effects culminated into a preserved aortic wall integrity in the absence of GPVI. Notably, pharmacological inhibition of GPVI using a specific blocking antibody further substantiated these protective effects in a preclinical setting, underscoring the therapeutic potential of selectively targeting platelet receptors in order to restrict AAA progression [16].

Given the critical role of platelets, particularly the platelet collagen receptor GPVI, in AAA progression, we aimed to investigate whether other platelet surface receptors may represent promising targets for pharmacological intervention in AAA. GPIbα, a key subunit of the GPIb–IX–V complex, mediates platelet adhesion to von Willebrand factor (VWF) and is indispensable for platelet recruitment under conditions of high shear stress. Beyond its classical role in thrombosis, GPIbα has been implicated in platelet-driven inflammatory responses and in vascular remodelling. However, experimental evidence defining the contribution of GPIbα-mediated platelet function and the underlying mechanisms governing AAA formation and progression remains limited [24]. Therefore, the present study investigated the role of GPIbα in AAA pathology using the IL-4Rα/GPIbα transgenic mice in a state-of-the-art *in vivo* model for experimentally induced AAA formation.

## 2. Methods

### 2.1 Study approval

All animal procedures were conducted in accordance to with internationally recognised ethical standards and relevant regulatory guideline. Experimental protocols were approved by the Ethics Committee of the State Ministry of Agriculture, Nutrition, and Forestry of North Rhine-Westphalia, Germany (permit IDs: 81-02.05.40.21.041, 81-02.04.40.2023.VG040, 81-02.04.2021.A429 and 81-02.04.2018.A409). Additional approval was granted by the Animal Care Committee of Heinrich Heine University. All experiments were performed in full compliance with the European Parliament’s Directive 2010/63/EU governing the protection of animals used for scientific research.

All studies involving human blood samples were conducted following written consent from participants in accordance with the requirements Ethics Committee of the University Hospital Düsseldorf, Germany (ATLANTA study, approval number: 2018-140-kFogU; biobank study, approval number: 5731R [2018-222_1]; MELENA study, approval number: 2018-248-FmB).

### 2.2 Animals

Specific pathogen-free IL-4Rα/GPIbα transgenic mice, in which the murine extracellular domain of GPIbα is replaced by the human IL-4 receptor were kindly provided by Prof. Dr. Carsten Deppermann (Centrum für Thrombose und Hämostase [CTH], Mainz, Germany). C57BL/6J mice were purchased from Janvier Labs. All experiments were exclusively conducted in male mice aged 10-12 weeks. Animals were maintained under standardised conditions in temperature-controlled housing (22 ± 1 °C) with a 12-h light–dark cycle. Mice were kept in type III Makrolon cages and provided unrestricted access to standard laboratory diet and water.

### 2.3 Human AAA samples

Fresh citrate-anticoagulated whole blood (BD–Vacutainer^®^; Becton, Dickinson and Company; #367714) was obtained in the department of Vascular- and Endovascular Surgery at the University Hospital Duesseldorf form patients diagnosed with AAA. Blood samples from healthy volunteers (average age: 65.5 ± 1.3) served as controls and were collected in the blood donor centre at the University Hospital of Duesseldorf, Germany.

### 2.4 Experimental AAA mouse model

For experimental induction of AAA formation in mice, the external porcine pancreatic elastase (ePPE) model was used. This approach involves the periadventitial application of porcine pancreatic elastase to the infrarenal abdominal aorta, thereby initiating aneurysm formation and enabling longitudinal *in vivo* assessment of disease progression. For surgical procedures, mice were anesthetised with 2–3% isoflurane. Preoperative analgesia was achieved by subcutaneous administration of buprenorphine (0.1 mg/kg body weight; Temgesic, Eumedica) 30 minutes prior to surgery. Anaesthesia was maintained and continuously monitored throughout the procedure. Anaesthesia was maintained and continuously monitored throughout the operation. The ePPE model was performed in accordance with previously established protocols described elsewhere [16, 19]. Briefly, the exposed infrarenal aorta was incubated with elastase (25.5 U/mL) for 5 minutes. Sham-operated animals underwent an identical procedure but received heat-inactivated elastase (100 °C for 30 minutes). Postoperative analgesia consisted of subcutaneous buprenorphine (0.1 mg/kg body weight) administered every 6 hours during the light phase. During the dark phase, buprenorphine was provided via drinking water (0.3 µg/mL) for three consecutive days. In addition, all mice received 0.1% β-aminopropionitrile (BAPN) in the drinking water continuously throughout the 28-day observation period, beginning 3 days before surgery [16]. For endpoint analysis and tissue harvesting, mice were euthanised on day 28 following ePPE induction by cervical dislocation under deep isoflurane anaesthesia.

### 2.5 Ultrasound imaging

AAA formation and progression in ePPE-treated mice were monitored by serial ultrasonography. The maximal luminal diameter of the infrarenal aorta within the aneurysmal segment was measured at baseline and on postoperative days 7, 14, 21, and 28. For imaging, mice were anaesthetised with 2–3% isoflurane and positioned on a temperature-controlled platform maintained at 37 °C. Ultrasound images were acquired using the Vevo 3100® High-Resolution In Vivo Micro-Imaging System (VisualSonics). Progressive aortic dilation was quantified using a standardised protocol based on longitudinal B-mode images captured during systole.

### 2.6 Blood Collection

Murine whole blood was collected into 300 µL of heparin solution (20 U/mL) for haematological analysis. Platelet, white blood cell (WBC) and red blood cell (RBC) counts as well as mean platelet volume (MPV), plateletcrit (PCT), platelet distribution width (PDW), haemoglobin (HGB) and haematocrit (HCT) were measured using an automated haematology analyser (Sysmex KX21N, Norderstedt).

### 2.7 Preparation of murine aortic tissue

At day 28 following ePPE surgery, mice were euthanised under deep isoflurane anaesthesia by cervical dislocation. Immediately thereafter, the thoracic cavity was opened, and the heart was accessed via puncture of the left ventricular apex with a butterfly needle to enable vascular perfusion. The vasculature was flushed with approximately 20 mL of cold heparin solution (20 U/mL, 4 °C, #2047217, Braun) to prevent coagulation. Subsequently, the abdominal aorta was carefully excised and fixed in 4% paraformaldehyde (#P087.5, Carl Roth) at 4 °C for 24 hours. Tissues were then dehydrated through a graded ethanol series, incubated in Roti®Histol (#6640.4, Carl Roth) for at least 12 hours at room temperature, and finally embedded in paraffin (#P3558, Sigma-Aldrich) for subsequent histological analysis.

### 2.8 Flow cytometry

Flow cytometry was employed to assess murine platelet function, glycoprotein expression, and platelet-leukocyte interactions. For murine studies, heparinised whole blood was washed three times with Tyrode’s buffer (137 mM NaCl, 2.8 mM KCl, 12 mM NaHCO₃, 0.4 mM NaH₂PO₄, 5.5 mM glucose; pH 6.5) via centrifugation at 650 × g for 5 min. The final cell suspension was resuspended in Tyrode’s buffer supplemented with 1 mM CaCl₂. To analyse platelet activation, washed platelets were incubated with fluorescently labelled antibodies against P-selectin (CD62P, Wug.E9-FITC, #D200, Emfret Analytics) and active integrin αIIbβ3 (JON/A-PE, #D200, Emfret Analytics) at a 1:10 dilution. Samples were stimulated with defined agonists, including adenosine diphosphate (ADP, #A2754, Sigma-Aldrich), collagen-related peptide (CRP, University of Cambridge, UK), thromboxane A2 analogue U46619 (#1932, Tocris), or PAR4 peptide (#3494, Tocris) for 15 min at 37 °C in the dark. Reactions were terminated by adding 300 µL of PBS. Platelet surface receptor profiling, washed blood was labelled with antibodies targeting integrin α5 (CD49e, Tap.A12-FITC, # M080-1, Emfret Analytics), integrin αIIbβ3 (CD41/61, Leo.F2-FITC, #M025-2, Emfret analytics), GPIX (CD42a, Xia.B4-FITC, #M051-1, Emfret analytics), GPIbα (CD42b, Xia.G5-PE, #M040-2), GPIbβ (CD42c, Xia.C3-FITC, #M050-1, Emfret analytics), GPV (CD42d, 1C2-APC, #148505, BioLegend), GPVI (JAQ1-FITC, #M011-1, Emfret analytics), FasL (CD178, MFL3-PE, 1:20, #106605, BioLegend), CD40L (TRAP/CD40L, MR1-FITC, #Ab24934, Abcam), or huIL4R (APC, #FAB230A, R&D Systems), at a 1:10 ratio for 15 min at room temperature. Upregulation of integrin α2 (CD49b, Sam.G4-FITC, #M070-1, Emfret Analytic) and integrin β3 (CD61, Luc.H11-FITC, #M031-1, Emfret Analytic) was measured following stimulation with the above agonists under the same conditions. For analysis of phosphatidylserine exposure, washed murine platelets were resuspended in binding buffer (10 µM HEPES, 140 µM NaCl, 2.5 mM CaCl_₂_; pH 7.4) and incubated with CyTM5 Annexin V (#559934, BD Biosciences) at a 1:10 dilution for 15 min at 37 °C in the dark in the presence of agonists. The reaction was stopped by adding 300 µL of binding buffer (0.1 M HEPES, 1.4 M NaCl, 25 mM CaCl_₂_; pH 7.4). To quantify platelet-leukocyte interactions, washed murine blood was labelled for 15 min at room temperature with antibodies against leukocytes (CD45, 30-F11-APC, BD Biosciences), neutrophils (Ly6G, 1A8-APC, BD Biosciences), T cells (CD3, 17A2-APC, BioLegend), B cells (CD19, 6D5-APC, #115512, BioLegend), macrophages/granulocytes (CD14, Sa14-2-APC, BioLegend and CD11b, M1/70-APC, BD Biosciences), with GPIX (CD42a, Xia.B4-FITC, #M051-1, Emfret analytics) as a platelet marker (all antibodies at a 1:10 dilution). In order to determine the formation of circulating platelet-RBC aggregates, whole blood samples were stained for RBCs (TER-119-FITC, 1:20, #11-5921-82, Invitrogen) and platelets (GPV, CD42d, 1C2-APC, #148505, BioLegend). To investigate VWF-binding, washed whole blood samples were labelled for VWF (VWF-FITC, 1:10, #P150-1, Emfret Analytics) and incubated with the indicated agonists. Samples were incubated for 15 min at RT in the dark and the reaction was stopped by adding 300 µL of PBS. All samples were analysed using a BD FACSymphony™ flow cytometer under standardised acquisition settings. For analysing human platelets, whole blood samples were diluted at a ratio of 1:10 in human Tyrode’s buffer (137 mM NaCl, 2.8 mM KCl, 12 mM NaHCO_3_, 0.4 mM NaH_2_PO_4_, and 5.5 mM glucose; pH 6.5) and specifically stained for 15 min at RT using an antibody against GPIbα (CD42b-FITC, #555472, BD Pharmingen^TM^) at a ratio of 1:10. All samples were analysed using a BD FACSCalibur™ flow cytometer.

### 2.9 Immunofluorescence staining of murine aortic tissue

Paraffin-embedded aortic segments from ePPE-treated mice were sectioned at 5 µm thickness using an automated microtome (Microm HM355, Thermo Fisher Scientific). Sections were deparaffinised, rehydrated, and subjected to heat-induced antigen retrieval in citrate buffer (pH 6.0) at 300 W for 10 minutes. For immunostaining, tissue sections were first blocked in DPBS containing 0.3% Triton X-100 and 5% goat serum for 1 hour at room temperature to prevent non-specific binding. Afterwards sections were incubated with a primary antibody for elastin (Eln, #ab307, Abcam, 1:50) overnight at 4 °C, followed by staining with Alexa Fluor™ Plus 555-conjugated secondary antibodies (#A32794, Invitrogen™, 1:200) for 1 hour at room temperature. Nuclei were counterstained with DAPI (#10236276001, Roche, 1:3000). Appropriate IgG controls were included for all stainings. Fluorescence images were generated using an LSM 880 Airyscan Fast system (Zeiss).

### 2.10 Histology of murine aortic tissue

Paraffin-embedded aortic segments from ePPE-treated mice were cut into 5 µm sections using an automated microtome (Microm HM355, Thermo Fisher Scientific). Haematoxylin and eosin (H/E) staining was performed following established protocols (27) to assess general tissue morphology. The thickness of the aortic media was quantified at ten evenly distributed locations within the aneurysmal segment using ZEN software (Zeiss, blue edition, version 3.3). Elastic fibre integrity was evaluated using modified Verhoeff–van Gieson (VVG) staining (#HT25A-1KT, Sigma-Aldrich) according to the manufacturer’s instructions. Elastin fragmentation was assessed using a graded classification system. Grade 1 indicated the absence of elastin fragmentation. Grade 2 was defined by involvement of up to 50% of the aortic wall. Grade 3 reflected fragmentation affecting more than 50% of the wall. Grade 4 denoted complete loss of intact elastic lamellae.

### 2.11 Enzyme-Linked Immunosorbent Assay (ELISA)

Plasma samples of AAA patients and controls from healthy volunteers and were analysed using a human glycocalicin ELISA kit (#MSB733858, MyBioSource). All ELISA were conducted according to the manufacturer’s instructions.

### 2.12 Turbidimetry

VWF concentration and activity in plasma samples of sham and ePPE-operated mice, as well as AAA patients and controls from healthy volunteers was analysed using turbidimetry as described elsewhere [25].

### 2.13 Statistical analysis

All data are presented as mean ± standard error of the mean (SEM). Statistical analyses were conducted using GraphPad Prism 8 (version 8.4.3). The sample size (*n*) indicated for each experiment represents the number of independent biological replicates. For murine experiments, statistical analyses were conducted under the assumption of normal data distribution, based on the use of inbred strains and the limited number of biological replicates for *in vivo* experiments. For human AAA datasets, normality was assessed using the Shapiro–Wilk test. Comparisons between two groups were performed using an unpaired multiple t-test, an unpaired student’s t-test or a Mann-Whitney U test. For correlation the Spearman correlation was used. Survival analyses were conducted using a Log-rank (Mantel-Cox) model. The AAA incidence in mice was analysed using a Fisher’s exact test. Statistical significance is denoted by asterisks (*p < 0.05; **p < 0.01; ***p < 0.001; ****p < 0.0001).

## 3. Results

### 3.1 Enhanced level of plasma glycocalicin in mice after experimentally induced AAA formation

In order to investigate the potential role of platelet GPIbα within AAA pathology, we initially employed the commonly used external porcine pancreatic elastase (ePPE) mouse model, which adequately reflects key pathological features of human AAA initiation and progression in mice. Aneurysm formation was longitudinally monitored over a time course of 28 days using high-resolution ultrasound imaging (**Figure 1A**). Thereby, ePPE-operated C57BL/6J mice revealed a significant increase in the inner aortic diameter as early as day 7 following surgery, with progressive aneurysmal expansion throughout the whole observation period (**Figure 1B**). In contrast, sham-operated mice receiving heat-inactivated elastase served as controls and consequently showed no evidence of aortic dilatation. Although the overall survival was moderately reduced within the group of ePPE-operated mice, this difference did not reach statistical significance compared with sham-operated controls (**Figure 1C**). Notably, the ePPE mouse model demonstrated an efficient and highly reproducible aneurysm induction, with all ePPE-operated mice exceeding the predefined cut-off threshold for AAA formation (≥150% increase in inner aortic diameter), whereas aneurysms were completely absent in sham-operated mice as indicated by AAA incidence (**Figure 1D**). Consistent with robust experimental AAA induction, severity classification at day 28 revealed that ePPE-operated mice predominantly developed moderate (200–250% increase in aortic diameter) or extremely severe aneurysms (≥400% increase), while sham-operated mice consistently showed no aneurysm formation (**Figure 1E** and **F**). To evaluate the relevance of platelet GPIbα in experimentally induced AAA, we first quantified platelet surface exposure of GPIbα at day 28 post-surgery in these mice via flow cytometry. Surface expression of GPIbα was comparable between both groups (**Figure 1G**). In contrast, plasma glycocalicin concentrations, reflecting proteolytic shedding of the GPIbα ectodomain, were significantly increased in ePPE-operated mice compared to sham controls (**Figure 1H**). Interestingly, plasma glycocalicin levels strongly correlated with progressive aortic diameter expansion, indicating that GPIbα shedding may be related to disease severity and aneurysm progression (**Figure 1I**). In contrast, analysis of the physiological GPIbα ligand von Willebrand factor (VWF) revealed no significant differences between sham and ePPE-operated mice, neither in plasma VWF activity nor in VWF antigen levels, indicating that experimental AAA is not associated with systemic alterations in VWF (**Figure 1K**–**M**). Collectively, these initial experimental data may identify platelet GPIbα ectodomain shedding as a previously unrecognised feature of experimental AAA and suggest plasma glycocalicin as a potential platelet-derived biomarker for disease monitoring and risk stratification.

**Figure 1.**
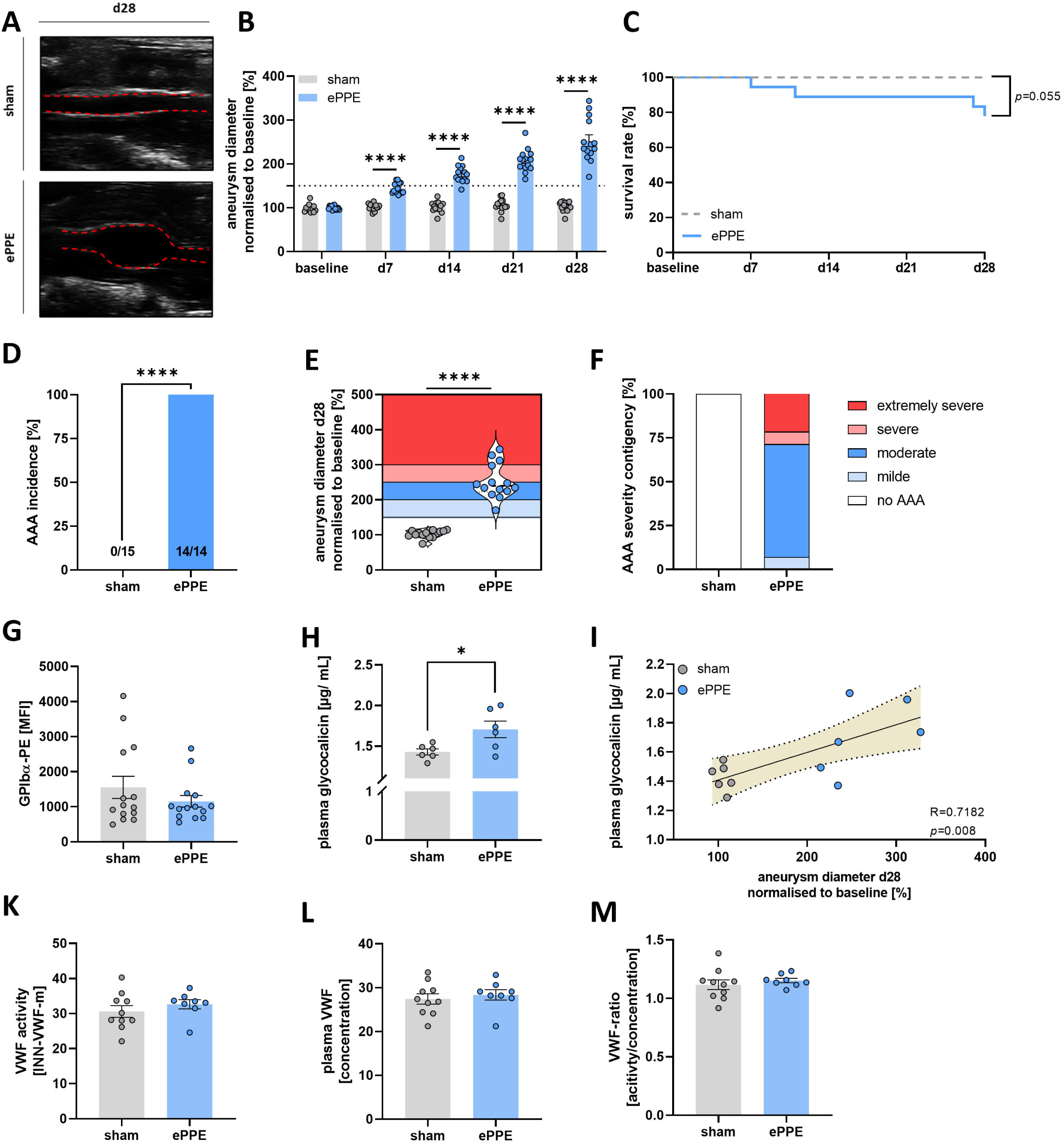
Enhanced level of plasma glycocalicin in mice after experimentally induced AAA formation. (**A**) Representative ultrasound images of sham and ePPE-operated C57BL/6J mice at day 28 post surgery (*n* = 14–15). (**B**) Aortic diameter progression of sham and ePPE-operated mice was tracked by ultrasound measurements over a time period of 28 days (*n* = 14–15). Data were normalised to baseline. (**C**) Survival rate of sham and ePPE-operated mice (*n* = 15–19) mice over a time period of 28 days. (**D**) AAA incidence of sham and ePPE-operated mice (*n* = 14–15) mice were defined by an aortic diameter ≥150% at day 28 post surgery. (**E** and **F**) AAA severity of sham and ePPE-operated mice (*n* = 14–15) mice at day 28 post surgery. (**G**) GPIbα surface exposure at the platelet surface of sham and ePPE-operated mice (*n* = 14) was analysed via flow cytometry. (**H**) Plasma glycocalicin concentrations of sham and ePPE-operated mice at day 28 post surgery (*n* = 6), analysed via ELISA. (**I**) Spearman’s correlation between plasma glycocalicin concentrations and aneurysm diameter at day 28 post ePPE surgery. (**K**–**M**) Plasma samples of sham and ePPE-operated mice were analysed for (**K**) VWF activity, (**L**) plasma VWF concentrations, and (**M**) the VWF activity/concentration ratio (*n* = 8–10) using turbidimetry. Data are represented as mean values ± SEM. Statistical analysis was performed using (**B**) a multiple t-test, (**C**) a Log-rank (Mantel-Cox) test, (**D**) a Fisher’s exact-test, and (**E**, **G**, **H** and **K**–**M**) an unpaired student’s t-test. ePPE = external porcine pancreatic elastase, MFI = mean fluorescence intensity, PE = phycoerythrin, VWF = Von Willebrand factor.

### 3.2 GPIb_α_ deficiency attenuates early aortic diameter expansion during experimentally induced AAA formation

In order to evaluate the functional contribution of platelet GPIbα to aneurysm progression, experimental AAA was induced in chimeric human interleukin-4 receptor (hIL-4R)/GPIbα-deficient mice (*Gp1ba^-/-^*), where the extracellular domain of human GPIbα was replaced by the corresponding extracellular domain of the human hIL-4Rα subunit [26]. Mice were analysed for AAA formation and progression using the ePPE mouse model for a time period of 28 days (**Figure 2A, Figure S1A** and **B**). Genetic deletion of GPIbα did not negatively affect the overall survival probability following experimental AAA induction compared to WT controls (**Figure 2B**). Longitudinal high-resolution ultrasound imaging was performed to assess aneurysm progression showing significantly attenuated early aneurysm expansion at day 7 post surgery (**Figure 2C**). In contrast, the aortic diameter at the observational endpoint (day 28) of AAA progression was comparable between WT and GPIbα-deficient mice suggesting a protective role of GPIbα only in the initial phase of AAA formation (**Figure 2D**). This was paralleled by an unaltered AAA incidence, with all ePPE-operated WT and GPIbα-deficient mice exceeding the predefined cut-off threshold for AAA formation (≥150% increase in inner aortic diameter) (**Figure 2E**). Consistently, aneurysm severity classification showed only moderate alterations between both groups (**Figure 2F** and **G**). In addition, GPIbα deficiency was associated with profound alterations in platelet-related haematological parameters. Loss of GPIbα resulted in reduced platelet counts and plateletcrit (PCT) (**Figure 2H** and **I**), accompanied by increased mean platelet volume (MPV) and platelet distribution width (PDW) (**Figure 2J** and **K**). However, these haematological alterations were also observed in naive (unoperated) GPIbα-deficient mice, indicating a genotype-dependent platelet phenotype rather than a consequence of the experimentally induced AAA formation (**Figure S1C**). Notably, WBC and RBC counts as well as haemoglobin (HGB) and haematocrit (HCT) were unaltered in both, naive and experimental AAA mice (**Figure S1C** and **Figure S2**).

**Figure 2.**
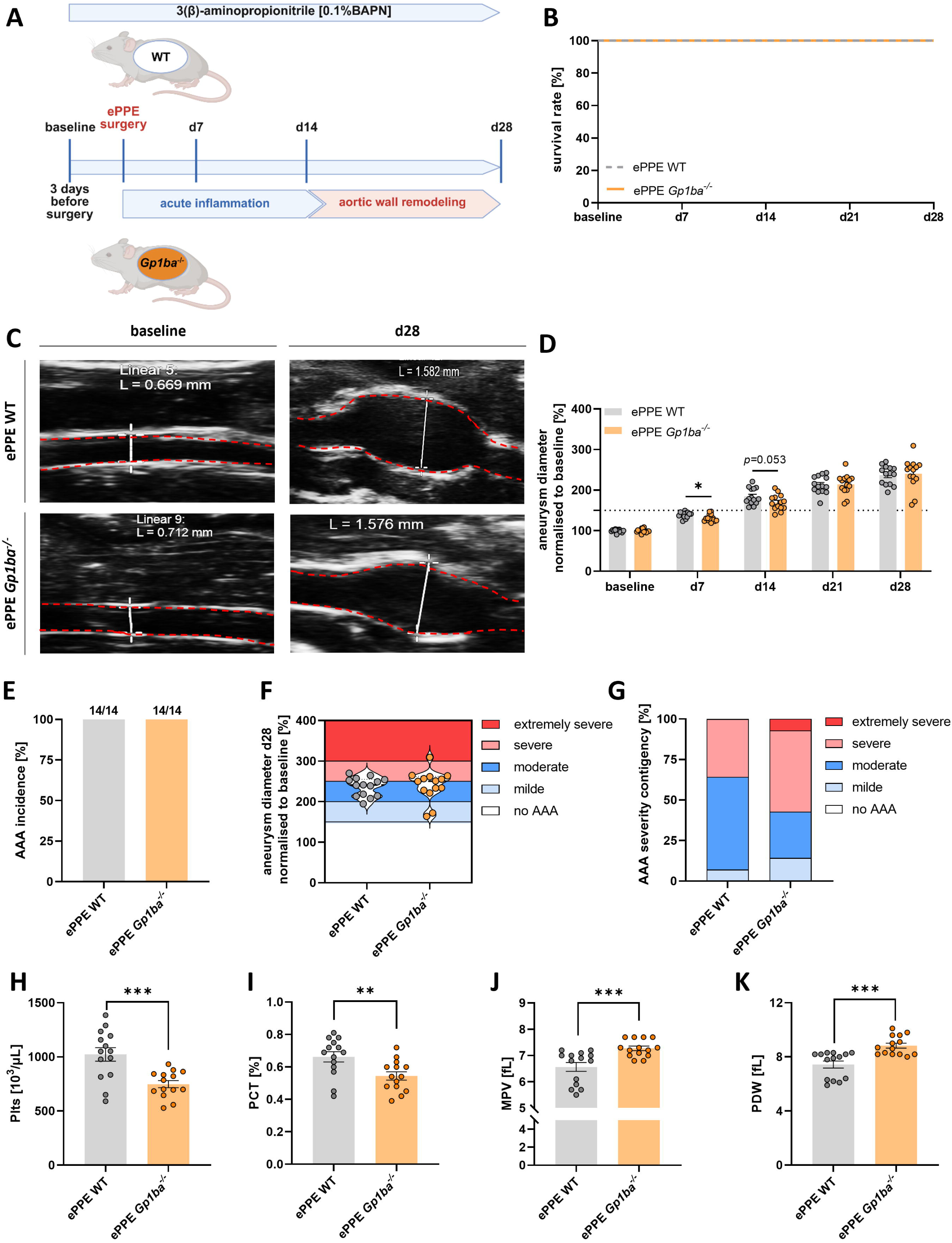
Genetic loss of GPIb_α_ attenuates early aortic diameter expansion in experimental abdominal aortic aneurysm (AAA) formation. (**A**) Schematic overview of the experimental procedure: AAA induction in WT and GPIbα-deficient (*Gp1ba*^-/-^) mice using the ePPE mouse model. Mice were sacrificed and analysed at day 28 post-surgery. (**B**) Survival rates of ePPE-operated WT and *Gp1ba*^-/-^ (*n* = 14) mice over a time period of 28 days. (**C**) Representative ultrasound images of ePPE-operated WT and *Gp1ba*^-/-^ (*n* = 14) mice before (baseline) and 28 days after surgery. (**D**) Aortic diameter progression in WT and *Gp1ba*^-/-^ (*n* = 14) ePPE mice was tracked by ultrasound measurements over a time period of 28 days. Data were normalised to baseline. (**E**) AAA incidence of ePPE operated WT and *Gp1ba*^-/-^ (*n* = 14) mice, defined by an aortic diameter ≥150% at day 28 post-surgery. (**F** and **G**) AAA severity of WT and *Gp1ba*^-/-^ (*n* = 14) mice at day 28 post ePPE surgery. (**H**–**K**) Platelet counts, PCT, MPV, and PDW of ePPE-operated WT and *Gp1ba*^-/-^ (*n* = 14) mice. Data are represented as mean values ± SEM. Statistical analysis was performed using (**B**) a Log-rank (Mantel-Cox) test, (**D**) a multiple t-test, (**E**) a Fisher’s exact-test and (**F, H**–**K**) an unpaired student’s t-test. BAPN = β-Aminopropionitrile, ePPE = external porcine pancreatic elastase, MPV mean platelet volume, PCT = plateletcrit, PDW = platelet distribution width, WT = wildtype.

### 3.3 GPIb_α_-deficient mice exhibit an attenuated aneurysm growth rate within early stage of AAA formation

To further investigate GPIbα−deficient mice in AAA, aneurysm growth rates were quantified by longitudinal ultrasound imaging throughout the 28-day observation period (**Figure 3A** and **B**). In line with reduced aortic diameter expansion at early time points, genetic deletion of platelet GPIbα significantly attenuated aneurysm growth rates during the initial phase of disease development at day 7 (**Figure 3A**). However, this early effect did not translate into a sustained long-term protection, as the overall aneurysm growth rate was unaltered between both groups (**Figure 3B**), resulting in the formation of aneurysmal tissue to a comparable extent (**Figure 3C**). As the progressive extracellular matrix degradation and elastin fragmentation are key pathological hallmarks of AAA which ultimately compromise aortic wall integrity, we next comprehensively analysed whether GPIbα deficiency affects the adverse structural remodelling of the aneurysmal vessel wall (**Figure 3D**–**F**). Histological analysis of H/E-stained aortic sections demonstrated no significant differences in aortic wall thickness between WT and GPIbα-deficient mice at day 28 following ePPE-induced AAA formation (**Figure 3D**). Consistently, elastin degradation grades were comparable between both genotypes (**Figure 3E**), culminating in an unaltered elastin content within the aneurysmal wall (**Figure 3F and Figure S3**). Collectively, these findings indicate that attenuated AAA formation at early time points can be compensated by other pathological mechanisms leading to comparable aneurysm diameter expansion at later time points in experimental mice with AAA.

**Figure 3.**
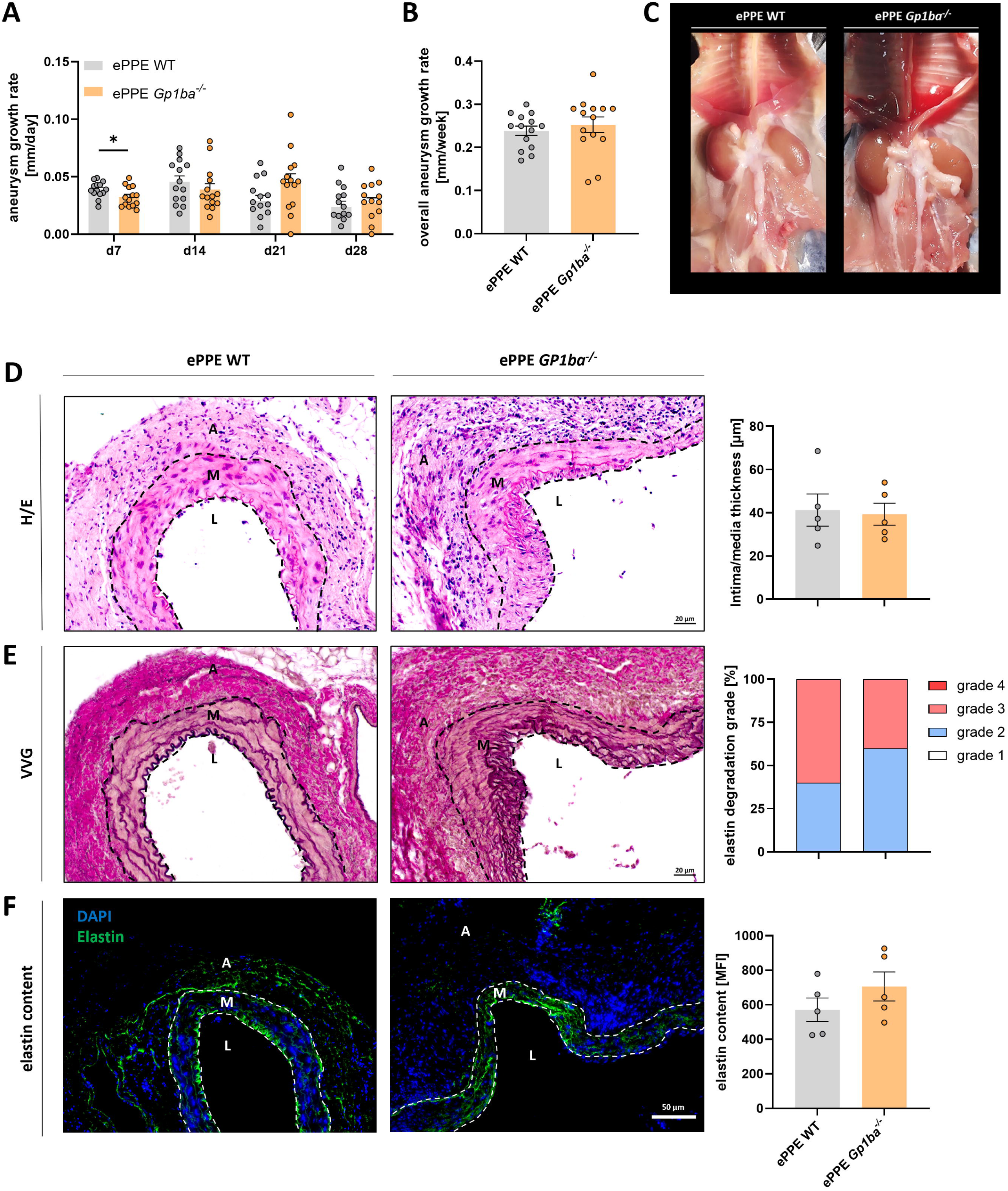
GPIb_α_-deficient mice exhibit an attenuated aneurysm growth rate within the early stage of AAA formation. **(A)** Aneurysm growth rate (mm/day) of ePPE-operated WT and *Gp1ba*^-/-^ (*n* = 14) mice over a time course of 28 days, as determined by ultrasound measurements. Data were normalised to baseline. (**B**) Overall aneurysm growth rate (mm/week) of WT and *Gp1ba*^-/-^(*n* = 14) ePPE mice. (**C**) Representative images of aortic tissue of ePPE-operated WT and *Gp1ba*^-/-^ (*n* = 14) mice at day 28 post surgery (**D**–**F**) Representative images and quantification of (**D**) the intima/media thickness (H/E staining), (**E**) elastin degradation (VVG staining. Grade 1: intact elastic lamellae, Grade 2: ≤ 50% elastin degradation within aortic tissue, Grade 3: ≥ 50% elastin degradation within aortic tissue, Grade 4: complete loss of elastic lamellae), and (**F**) elastin content (anti-Eln/Cy3; green) of aortic tissue of ePPE-operated WT and *Gp1ba*^-/-^ (*n* = 5) mice at day 28 post-surgery. Nuclei were stained with DAPI (blue). Data are represented as mean values ± SEM. Statistical analysis was performed using (**A**) a multiple t-test and (**B, D**–**F**) an unpaired student’s t-test. A = Adventitia, ePPE = external porcine pancreatic elastase, H/E = hematoxylin/eosin, L = Lumen, M =Media, VVG = Verhoeff-van Gieson, WT = wildtype.

### 3.4 GPIb_α_ deficiency results in pronounced platelet activation upon experimental induced AAA formation

To determine whether genetic GPIbα deletion intrinsically alters platelet function in the transgenic mouse model used herein, we performed an initial and comprehensive phenotypic characterisation of platelets derived from naive GPIbα-deficient mice (**Figure S4**). Under naive conditions, GPIbα-deficient platelets exhibited a constitutively hyperreactive phenotype as characterised by increased integrin α_IIb_β_3_ activation (JON/A) and elevated P-selectin exposure, reflecting α-granule secretion. This hyperactive state was detected under resting conditions and following stimulation of G-protein-coupled receptors using ADP or the PAR4 agonist peptide (only integrin activation) at low and high concentrations (**Figure S4A.** In addition, various receptors at the platelet surface of GPIbα-deficient mice, including integrin α_2_, integrin α_IIb_β_3_, and GPV, were significantly upregulated (**Figure S4B, E** and **F**). In contrast, platelet surface expression of CD40L and Fas ligand (FasL), as well as phosphatidylserine (PS) exposure assessed by Annexin V binding, remained unaltered, indicating that neither platelet inflammatory signalling nor procoagulant activity was intrinsically affected by the loss of GPIbα in the here used mouse model (**Figure S4C**, **D** and **H**). However, genetic GPIbα deletion significantly reduced the binding of its physiological ligand VWF to the platelet surface, confirming the functional loss of the GPIbα–VWF axis (**Figure S5**). Following ePPE-induced AAA formation, GPIbα-deficient mice exhibited an even more pronounced hyperreactive platelet phenotype. At day 28 after surgery, platelets showed a persistently elevated basal activation profile, as evidenced by increased degranulation (P-selectin exposure) and integrin α_IIb_β_3_ activation (JON/A). under resting conditions (**Figure 4A**). Moreover, agonist-induced platelet activation was markedly enhanced following G-protein coupled receptor activation, as observed already in naive mice. However, GPVI-mediated activation of platelets using collagen-related peptide (CRP), led to significantly elevated integrin α_IIb_β_3_ activation in GPIbα-deficient mice at day 28 post surgery demonstrating a broad increase in platelet responsiveness in experimental AAA in transgenic mice (**Figure 4A**). Phenotypic analysis further revealed enhanced surface expression of multiple platelet adhesion receptors, including integrin α_2_, integrin α_IIb_β_3_, integrin β_3_, integrin α_5_, GPV, and GPIX, following experimental AAA induction in GPIbα-deficient mice as already observed in naive mice (**Figure 4B, E** and **F**). In contrast to naive mice, we detected elevated platelet surface expression of the pro-inflammatory mediator CD40L and the prothrombotic FasL in GPIbα-deficient mice following experimental AAA induction (**Figure 4C** and **D**). In line with these pronounced activation-associated alterations, PS exposure of platelets was significantly enhanced after experimental AAA induction as well (**Figure 4H**). Notably, this enhanced procoagulant activity in transgenic mice was not observed under naive conditions, indicating an AAA induced phenotype (**Figure S4H**). In addition, the formation of circulating platelet-leukocyte and platelet-RBC aggregates during AAA formation was unaffected by GPIbα deficiency (**Figure 4I** and **J**). These findings demonstrate that the here used transgenic mouse model with GPIbα deficiency induces a constitutively hyperreactive platelet phenotype, which is further amplified during experimental AAA, accompanied by a pronounced prothrombotic phenotype.

**Figure 4.**
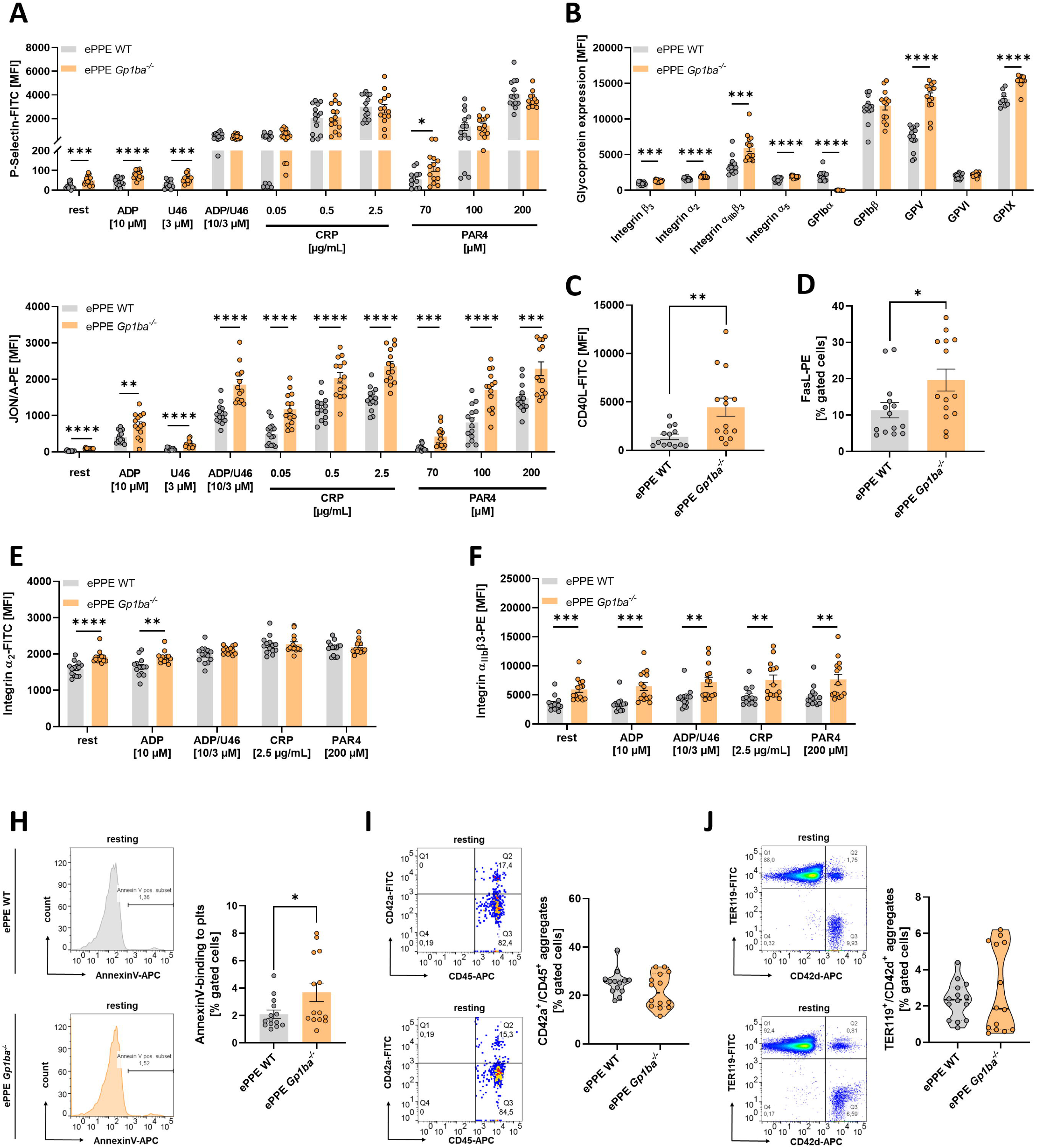
GPIb_α_ deficient mice show pronounced platelet hyperactivity in experimental AAA. (**A**–**I**) Washed murine whole blood of ePPE-operated WT and *Gp1ba*^-/-^ mice was analysed by flow cytometry at day 28 post-surgery. (**A**) Platelet degranulation (P-selectin-FITC) and integrin α_IIb_β_3_ activation (JON/A-PE) (*n* = 13), (**B**) platelet surface exposure of integrin β_3_, integrin α_2_, integrin α_IIb_β_3_, integrin α_5_, GPIbα, GPIbβ, GPV and GPVI were analysed (*n* = 9–14). Platelets were stimulated with indicated agonists. (**C** and **D**) Surface externalisation of (**C**) CD40L (*n* = 13–14) and (**D**) FasL (*n* = 14) on the platelet surface under resting conditions. (**E** and **F**) Externalisation of (**E**) integrin α_2_(*n* = 14) and (**F**) integrin α_IIb_β_3_(*n* = 14) on the platelet surface. Platelets were stimulated with indicated agonists. (**H**) PS exposure of platelets was analysed via Annexin V binding to platelets (*n* = 14). (**I** and **J**) Aggregate formation of platelets with (**C**) leukocytes (*n* = 14) and (**D**) RBCs (*n* = 14) was analysed as double-positive events for CD42d (platelet maker, GPV) and CD45 (leukocyte marker), respectively TER119 (RBC marker). Data are presented as mean ± SEM. Statistical analyses was performed using (**A**, **B**, **E** and **F**) a multiple t-test and (**C**, **D, H**–**J**) an unpaired student’s t-test. ADP = Adenosine diphosphate, APC =allophycocyanin, CRP = collagen-related peptide, ePPE = external porcine pancreatic elastase, FITC = fluorescein isothiocyanate, GP = glycoprotein, MFI = mean fluorescence intensity, PAR4 = protease-activated receptor 4 activating peptide, PE = phycoerythrin, Plts = platelets, U46 = thromboxane A2 analogue U46619, WT = wildtype.

### 3.5 Platelets derived from AAA patients display increased GPIb_α_ surface expression accompanied by elevated plasma VWF activity

Given the emerging role of the intraluminal thrombus (ILT) as a dynamic inflammatory neo-tissue in human AAA, we next investigated the impact of platelet GPIbα and its ligand VWF to ILT formation and structure. The ILT is continuously renewed through its interaction with circulating blood and constitutes a highly proteolytic and oxidative microenvironment that promotes progressive degradation of the underlying aortic wall [27, 28]. Histologically, the ILT exhibits a characteristic multilayered architecture composed of distinct luminal, medial, and abluminal regions [29]. To determine the spatial distribution of platelet-derived components within the human ILT, immunohistochemical staining for GPIbα and VWF was performed on sections from human AAA specimens (**Figure 5A** and **B**). Baseline clinical characteristics of the AAA patient cohort included in this study are summarised in **Table 1**. Spatial analysis revealed a highly compartmentalised distribution of GPIbα within the ILT, with marked enrichment in the luminal layer, whereas GPIbα was moderately present in the medial and abluminal regions (**Figure 5A**). A comparable distribution pattern was observed for VWF, which was likewise highly enriched in the luminal layer (**Figure 5B**). High-magnification images confirmed the preferential localisation of both GPIbα and VWF to the luminal layer, indicating an accumulation of platelet-rich thrombotic material at the blood-exposed interface of the ILT (**Figure 5A** and **B**). This was highlighted by a quantitative analysis, which demonstrated a significant enrichment of both GPIbα- and VWF-positive areas within the luminal layer compared with the medial/abluminal region of the ILT (**Figure 5C** and **D**). Quantitative assessment of the overall ILT composition of human AAA explants revealed that platelets and VWF accounted for 11.32% and 18.46% of the total ILT tissue, respectively, whereas the remaining 70.22% was composed of other structural components such as fibrinogen (**Figure 5E**). Notably, these findings identify a pronounced spatial compartmentalisation of platelets and VWF within the human ILT, thus supporting the concept of an ongoing platelet recruitment – likely facilitated via VWF – at the blood-ILT interface, which may directly contribute to continuous thrombus renewal within AAA pathology. To determine whether platelet GPIbα and its physiological ligand VWF are systemically altered in human AAA, circulating levels of VWF and GPIbα-associated biomarkers were quantified in samples obtained from patients with AAA and healthy controls (**Figure 5F**–**J**). In this context, circulating VWF antigen levels were comparable between patients with AAA and healthy controls, indicating that total plasma VWF abundance remained unaltered (**Figure 5F**). However, VWF activity was significantly increased in patients with AAA (**Figure 5G**), culminating into a markedly shifted VWF antigen-to-activity ratio, thus indicating enhanced functional VWF activity in the AAA cohort (**Figure 5H**). In parallel, platelets from patients with AAA exhibited significantly increased surface expression of GPIbα compared with healthy controls (**Figure 5I**). Despite this enhanced platelet GPIbα expression, plasma glycocalicin levels remained unchanged, indicating normal GPIbα ectodomain shedding (**Figure 5J**). Collectively, these findings demonstrate enhanced platelet GPIbα surface expression and increased activity of circulating VWF, suggesting an important role of the GPIbα–VWF axis in human AAA pathology.

**Figure 5.**
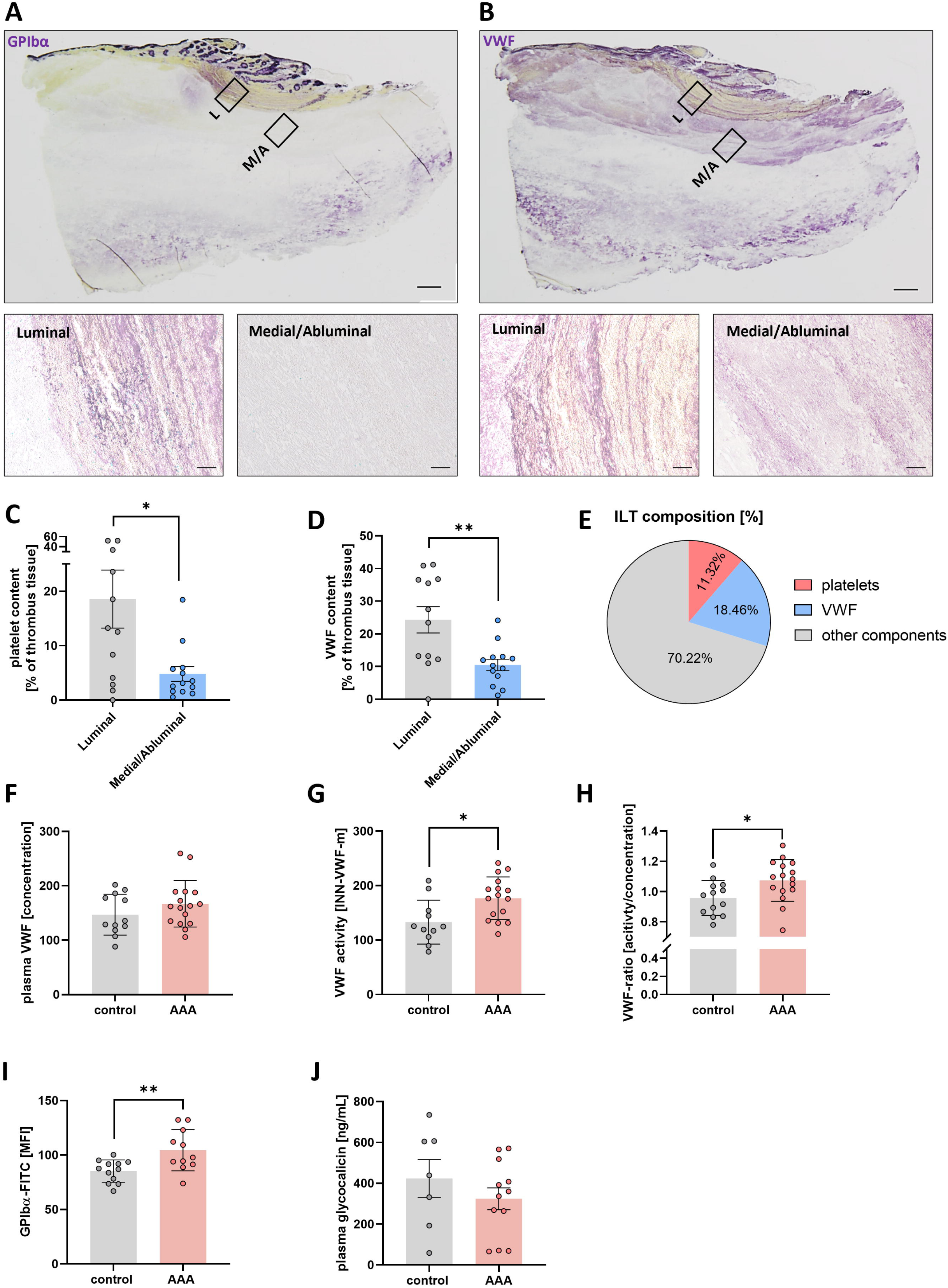
Platelets derived from AAA patients display increased GPIb_α_ surface expression accompanied by elevated plasma VWF activity. (**A**–**E**) Representative images and quantification of ILT tissue samples from AAA patients (*n* = 13). Thrombus sections were immunohistochemically stained and quantified for (**A** and **C**) platelets (anti-GPIbα) and (**B** and **D**) VWF (anti-VWF). Scale bars: 500 µm (overview) and 50 µm. (**E**). Relative composition of human ILT tissue samples with regard to platelet and VWF content (*n* = 13). (**F**–**H**) Plasma samples of AAA patients and healthy volunteers were analysed for (**F**) plasma VWF concentration, (**G**) VWF activity, and (**H**) the VWF activity/concentration ratio (*n* = 11–16) using turbidimetry. (**I**) GPIbα surface exposure at the surface of platelets from healthy volunteers and AAA patients (*n* = 11–16) was analysed via flow cytometry. (**J**) Plasma glycocalicin concentrations of controls from healthy volunteers and AAA patients were analysed by ELISA (*n* = 7–12). Data are represented as mean values ± SEM. Statistical analysis was performed using (**C** and **D**, **F**–**J**) a Mann-Whitney U test. FITC = fluorescein isothiocyanate, ePPE = external porcine pancreatic elastase, ILT = Intraluminal thrombus, L = Luminal, M/A= Medial/Abluminasl, MFI = mean fluorescence intensity, PE = phycoerythrin, VWF = Von Willebrand factor.

## 4. Discussion

Beyond their canonical function in haemostasis, platelets are increasingly recognised as active regulators of cardiovascular pathologies. Dysregulated platelet activation and functional reprogramming emerged as prompt drivers of a broad range of cardiovascular disorders, including ischemic stroke, atherosclerosis, and myocardial infarction [14, 30–33]. In this context, accumulating experimental and clinical evidence indicates a central role of sustained platelet hyperactivation in AAA pathogenesis. This may reinforce thrombo-inflammatory processes within the aneurysmal wall and the ILT, directly contributing to AAA initiation and progression [8, 19, 34, 35]. These findings highlight platelet-specific signalling pathways as promising and novel therapeutic targets to effectively restrict aneurysm growth. Consistently, multiple preclinical studies have demonstrated beneficial effects of antiplatelet therapies that significantly attenuate aortic diameter expansion, as well as rupture risk across distinct experimental AAA models [36]. In parallel, several experimental studies identified novel platelet-specific targets that were shown to effectively restrict aneurysm growth in preclinical settings, including GPVI, olfactory receptor 13 (Olfr13), as well as the CD36–thrombospondin-1 (TSP-1) axis [16, 19, 21, 23]. However, successful translation of these promising preclinical findings into clinical practice remains challenging, underscoring the relevance to identify novel, pharmacological targets.

The GPIb–IX–V complex serves as a critical modulator of platelet recruitment to the site of the injured vessel wall under high arterial shear conditions by mediating the interaction between platelet GPIbα and immobilised or shear-activated VWF [37, 38]. This transient ligand–receptor interaction enables platelet tethering and adhesion, as well as initiates intracellular signalling that promotes subsequent activation of the integrin α_IIb_β_3_. Engagement of activated integrin α_IIb_β_3_ with fibrinogen subsequently stabilizes platelet adhesion to the ECM and facilitates platelet–platelet interactions, thereby promoting thrombus growth [39–41]. Beyond its role in the initial stage of platelet adhesion, the GPIbα–VWF axis also contributes to thrombus growth via interactions with VWF presented on activated platelets within the developing thrombus [42, 43]. Moreover, GPIb–IX–V signalling is capable of reinforcing platelet procoagulant responses, thus promoting local thrombin generation and fibrin formation. Collectively, the GPIbα–VWF axis represents a key driver of platelet recruitment, activation, and thrombus maturation, particularly under high shear conditions as found in the abdominal aorta [43–45]. Accordingly, GPIbα-dependent platelet adhesion represents a critical initiating mechanism in arterial thrombus formation and has been implicated in the pathogenesis of major thrombotic disorders, including myocardial infarction and ischemic stroke [46–49]. In this context, multiple studies have reported a distinct prothrombotic platelet phenotype in patients with AAA. Circulating levels of soluble P-selectin were markedly elevated, indicating increased platelet activation *in vivo* [50]. Moreover, platelets isolated from AAA patients exhibit a persistent hyperactive and -reactive phenotype, characterised by enhanced P-selectin surface exposure and increased α_IIb_β_3_ integrin activation, both under basal conditions and following agonist stimulation [8]. In parallel, increased platelet procoagulant activity, as well as the formation of a platelet-rich ILT, further supporting a sustained platelet-driven prothrombotic state in AAA [8, 51]. Collectively, these observations position the GPIbα–VWF axis as a potential mediator of thrombo-inflammatory mechanisms underlying AAA initiation and progression. However, the extent to which GPIbα-dependent platelet recruitment and adhesion contribute to AAA pathology remains largely unknown.

In the present study we identified platelet GPIbα as a mediator of early aneurysm development. Genetic ablation of GPIbα significantly attenuated early aortic expansion in the ePPE model for experimental *in vivo* AAA induction, indicating a critical role for GPIbα-dependent platelet function during the initial stages of AAA. However, attenuated early aortic expansion in the ePPE model of experimental AAA was restricted to the initial phase of aneurysm formation (day 7 post-surgery) (**Figure 2D**). The transient nature of this effect may support the concept that platelet recruitment constitutes a critical early event in the inflammatory response underlying aneurysm initiation. In this context, several experimental studies across complementary murine *in vivo* AAA models, namely the AngII, PPE, and ePPE models, have consistently demonstrated a pronounced recruitment of both platelets and inflammatory leukocytes as a characteristic feature of the inflammatory response in early experimental AAA formation [16, 20, 52–54]. Given the central role of the GPIb–IX–V complex as crucial mediator of platelet tethering and recruitment under high shear – as apparent within the abdominal aorta – loss of GPIbα may substantially restrict platelet recruitment at the site of aneurysm formation, thus attenuating platelet-mediated local inflammation. Preclinical studies have reported that neutrophil recruitment and migration into the aneurysmal segment is at least partially regulated in a platelet-dependent manner [16, 20]. The pathological relevance of platelet-neutrophil interactions is further supported by Eliason *et al.* who demonstrated compelling evidence, that neutrophil-driven inflammation is a critical driver of aneurysm progression, as antibody-mediated neutrophil depletion substantially attenuates aortic expansion in experimental AAA [55]. Notably Corken *et al.* demonstrated a GPIb-IX–dependent role in immune regulation under systemic inflammatory conditions using the same hIL-4R/GPIbα-deficient mouse model employed in the present study. Thereby, genetic deletion of GPIbα markedly restricted platelet–neutrophil and – monocyte interactions not only under basal conditions but also following induction of systemic inflammation by cecal ligation and puncture (CLP) [56]. This observation, however, is not surprising especially giving the established role of the GPIbα–Mac-1 axis as a crucial molecular mechanism for platelet–leukocyte interactions, thus mediating platelet–immune cell crosstalk under inflammatory conditions [57, 58]. Attenuation of this GPIbα-mediated platelet–neutrophil interactions may therefore contribute – at least in part – to the transient reduction in aneurysm expansion observed in the present study, while underscoring the stage-specific contribution of GPIbα to AAA pathogenesis. Despite significantly affecting early aneurysm growth, the protective effect of GPIbα deficiency was compensated during subsequent AAA progression. In line with this, aortic wall integrity within the aneurysm segment – as indicated by elastin fragmentation – was unaffected by GPIbα deficiency. Consequently, neither AAA incidence nor overall aneurysm severity revealed major differences, indicating that platelet GPIbα may primarily contributes to aneurysm initiation rather than sustained disease progression. In this context, the lack of sustained aortic diameter restriction may be attributable to the pronounced and persistent platelet hyperactivity observed in GPIbα-deficient mice (hIL-4R/GPIbα). As previously stated, platelets derived from these mice already display a prothrombotic phenotype under naive conditions (**Figure S3**), which is further amplified following experimental AAA induction (**Figure 4**). Consistent with our findings, Deng *et al.* previously reported a constitutively hyperreactive platelet phenotype in hIL-4R/GPIbα-deficient mice, characterised by enhanced intracellular calcium mobilisation and increased P-selectin surface exposure [59]. Mechanistically, this pronounced hyperactive phenotype has been attributed to the unfolded GPIbα trigger sequence retained within the chimeric IL-4Rα extracellular domain, thereby constitutively activating intracellular GPIb–IX downstream signalling pathways, resulting in persistent platelet activation and a prothrombotic phenotype [26, 59]. From a mechanistic perspective, binding of multimeric VWF to platelet GPIb–IX–V serves as a critical molecular linker between shear-dependent platelet adhesion and intracellular activation. Receptor engagement initiates activation of tyrosine kinases, including Syk, FAK, and Pyk2, followed by downstream PLC, PLA_2_, and PI3K signalling. These signalling cascades ultimately result in integrin α_IIb_β_3_-dependent platelet activation [41, 43, 60, 61]. In the context of AAA, Wagenhäuser *et al.* recently reported sustained platelet hyperactivation (integrin α_IIb_β_3_ activation and P-selectin exposure) as a pathological feature of AAA progression in both mice and humans. In the PPE model, platelet activation progressively increased during AAA progression, indicating that aneurysm growth is capable of dynamically altering the platelet activation profile in a time-dependent manner [8]. Consistent with these observations, Morrell *et al.* demonstrated enhanced platelet degranulation, reflected by increased P-selectin exposure, in the ePPE mouse model [21]. Collectively, these findings support the concept that AAA progression is accompanied by progressive platelet priming and hyperactivity. However, this disease-related amplification of platelet activation may provide a mechanistic basis for the pronounced hyperactive platelet phenotype observed in GPIbα-deficient mice following experimental AAA in the present study (**Figure 4**). Given the established role of platelet activation in AAA pathogenesis, the pronounced hyperactive phenotype (**Figure 4A**) and enhanced surface receptor expression (**Figure 4B**) observed in GPIbα-deficient mice may progressively offset the initial protective effect conferred by genetic loss of GPIbα. Moreover, GPIbα deficiency was associated with pronounced platelet PS exposure, particularly following experimental AAA induction, indicating a substantially enhanced procoagulant activity. Importantly, different studies demonstrated that procoagulant activity of platelet is markedly increased during AAA development in both murine AAA models and in patients diagnosed with AAA [8, 19]. In experimental AAA, platelet PS exposure positively correlates with diameter expansion in the PPE model, further implicating procoagulant activity as a platelet-specific feature of AAA pathology [8]. Collectively, the pronounced prothrombotic phenotype in GPIbα-deficient mice upon experimental AAA induction may – at least in part – account for the subsequent convergence in aneurysm progression observed at day 28 (**Figure 2D**), suggesting that platelet GPIbα primarily promotes aneurysm initiation rather than sustained disease progression.

To assess the clinical relevance of the GPIbα-VWF axis for human AAA pathology, spatial profiling of human ILT specimens from patients with AAA revealed a highly compartmentalised distribution of GPIbα and VWF, with pronounced enrichment within the luminal ILT layer. Especially VWF was highly abundant within human ILT tissue, accounting for approximately 18% of the total thrombus composition. In this context, Touat *et al*. already demonstrated that continuous luminal remodelling of the ILT is mainly driven by its persistent exposure to circulating blood. This ongoing thrombus renewal is accompanied by pronounced local platelet activation and procoagulant activity, particularly within the luminal thrombus layer, where platelet-derived markers including sP-selectin, sCD40L, and sGPV were found to be markedly enriched. Importantly, these thrombus-associated mediators were also elevated in the circulation of patients with AAA [50]. These findings suggest that ongoing platelet activation and recruitment at the blood–thrombus interface contribute to a systemic prothrombotic state, thus directly modulating the luminal renewal of the ILT. However, the here shown spatial distribution of both GPIbα and VWF within the luminal layer further strengthens the concept of ongoing platelet recruitment to the ILT – likely facilitated via VWF – at the blood-ILT interface. These findings were further supported by recent proteomic profiling of human ILT specimens using liquid chromatography in combination with mass spectrometry, which identified a significantly enhanced abundance of VWF within the ILT compared with thrombi generated from the blood of healthy donors. Notably, the GPIbβ subunit of the GPIb–IX–V complex was found to be significantly increased within the ILT tissue, whereas the abundance of GPIbα remained unaltered [35]. Collectively, these findings strongly implicate the GPIbα–VWF axis as a key mediator of persistent platelet recruitment to the aneurysmal site, potentially driving AAA progression by promoting both, ILT renewal and growth. Thus, GPIbα–VWF driven thrombus remodelling in AAA might serve as a potential novel therapeutic target.

Beyond these findings, patients with AAA exhibited significantly elevated circulating VWF activity, resulting in a markedly reduced VWF antigen-to-activity ratio indicative of a shift towards a more functionally active VWF phenotype. Importantly, we further demonstrated for the first time that circulating platelets from AAA patients display significantly increased surface expression of GPIbα compared to healthy controls (**Figure 5**). Over the past years, several platelet-derived parameters have emerged as potential biomarkers for the prediction and risk stratification in AAA. In particular, elevated circulating levels of sGPVI have consistently been reported to be associated with AAA formation and disease progression [16, 23]. Importantly, Benson *et al.* independently reproduced this linkage in two geographically distinct patient cohorts, strengthening the translational relevance of sGPVI as a circulating biomarker. Notably, sGPVI demonstrated a stronger association with aneurysm growth than D-dimer, currently one of the most widely established circulating biomarkers of AAA [23]. These findings highlight the potential of platelet-specific biomarkers to improve the detection and risk stratification of patients with AAA and support their significance as clinically applicable indicators of disease progression. Accordingly, the identification and successful clinical implementation of novel platelet-specific biomarkers remain highly relevant for improving AAA diagnosis, disease monitoring, and risk stratification. In this context, the markedly enhanced platelet surface expression of GPIbα observed in the present study provides first preclinical evidence supporting GPIbα as a potentially biomarker for AAA. However, further longitudinal studies in larger, well-characterised patient cohorts will be required to validate the diagnostic and prognostic potential of platelet GPIbα as a robust and effective predictor of AAA disease progression and to establish its utility for risk stratification.

Collectively, this study identified a dysregulated GPIbα–VWF axis in human AAA pathology, mainly characterised by enhanced platelet GPIbα surface expression and markedly increased activity of circulating VWF. These findings provide first preclinical evidence that platelet GPIbα may potentially serve as biomarker for AAA patients. In addition, the pronounced spatial distribution of both GPIbα and VWF within the luminal ILT layer further supports the concept of sustained platelet recruitment to the ILT at the blood–ILT interface, probably driven by VWF-mediated platelet adhesion. This persistent platelet recruitment may contribute to continuous ILT renewal and growth, thus sustaining the thrombo-inflammatory environment to reinforce the progression of AAA.

## Nonstandard Abbreviations and Acronyms

AAA: abdominal aortic aneurysm
BAPN: β-aminopropionitrile
CRP: collagen-related peptide
GP: glycoprotein
ILT: intraluminal thrombus
MMP: matrix metalloproteinases
MPV: mean platelet volume
PAR4: protease-activated receptor 4
ePPE: external pancreatic porcine elastase
PS: phosphatidylserine
U46: thromboxane A2 analogue U46619
VVG: Verhoeff-van Gieson
WT: wildtype

## Acknowledgments

We thank Martina Spelleken, Michaela Steffen Fritzke and Doga Sahin for excellent technical assistance and Carsten Deppermann for providing IL-4Rα/GPIbα transgenic mice. We would like to acknowledge the Centre for Advanced Imaging (CAi) at Heinrich-Heine-University Düsseldorf for providing access to the Zeiss LSM 880 Airyscan Fast system (DFG-INST 208/746-1 FUGG) and especially for providing support during imaging and analysis. All graphical abstracts were generated with BioRender.com.

## Author Contributions

ME and KJK designed the study. AB, KJK, MC, JO, AS, and DS performed experiments. AB, KJK, ME analysed and interpreted the data. CD provided hIL4Rα/GPIbα mice. MUW and HS provided human AAA samples. AB, KJK and ME wrote the manuscript with all authors providing feedback. KJK and ME contributed equally to this study.

## Data Availability

The original contributions presented in the study are included in the article. Further inquiries can be directed to the corresponding author.

## Supplemental Material

Figure S1-S5

