## Supplemental data for "Dysregulated Platelet GPIbα–VWF Signalling in Abdominal Aortic Aneurysm formation and Progression"

### **Supplemental Material**

#### **Dysregulated platelet GPIIb/IIIa–VWF signalling contributes to Abdominal Aortic Aneurysm formation and Progression**

Feige T<sup>1\*</sup>, Krott KJ<sup>1\*</sup>, Bosbach A<sup>1</sup>, Chario M<sup>1</sup>, Ortscheid J<sup>1</sup>, Saleem A<sup>1</sup>, Wagenhäuser MU<sup>1</sup>, Schelzig H<sup>1</sup>, Elvers M<sup>§1</sup>

\*Contributed equally

§ corresponding author

<sup>1</sup>Department of Vascular- and Endovascular Surgery, University Hospital Duesseldorf, Heinrich-Heine University, Duesseldorf, Germany

### Supplemental figures

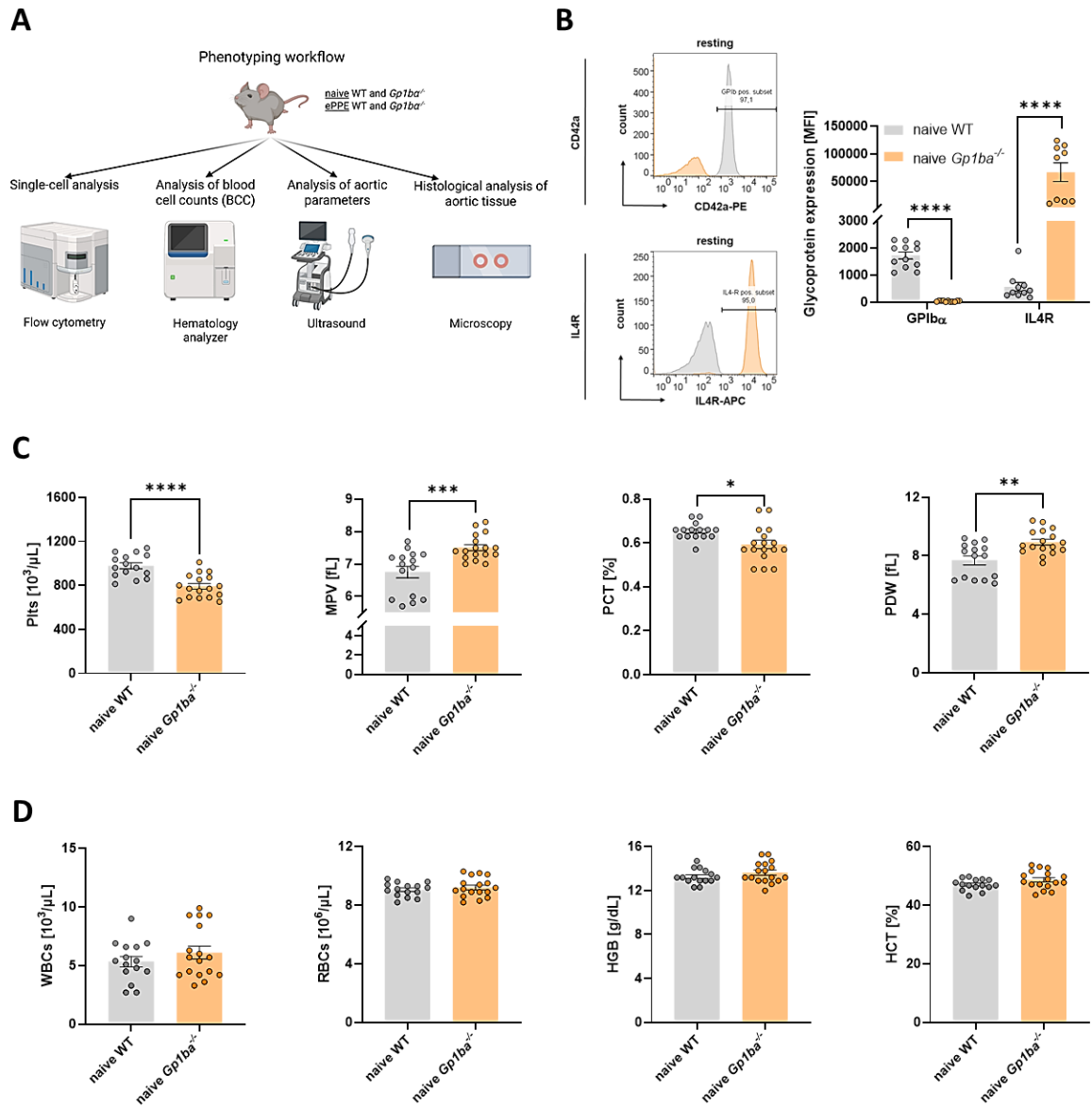

**Figure S1. Genetic loss of GPIIbα in mice alters platelet counts and phenotypic characteristics.**

(A) Schematic overview of phenotyping workflow of naïve and ePPE operated WT (*Gp1ba*<sup>+/+</sup>) and GPIIbα-deficient (*Gp1ba*<sup>-/-</sup>) mice. Ultrasound measurements were performed to analyse aortic parameters. Flow cytometry was used for single-cell analysis of whole blood samples. Blood cell counts were quantified by Sysmex haematology analyser, and aortic tissue sections were prepared for histological analysis. (B) For genotyping of mice, washed whole blood of naïve WT and *Gp1ba*<sup>-/-</sup> mice was analysed for platelet surface exposure of GPIIbα ($n = 11$–$12$) and IL4R ($n = 9$). (C) Platelet counts, MPV, PCT and PDW of naïve WT ($n = 15$) and *Gp1ba*<sup>-/-</sup> ($n = 17$) mice. (D) WBC and RBC counts, respectively HGB and HCT of naïve WT ($n = 15$) and *Gp1ba*<sup>-/-</sup> ($n = 17$) mice. Data are presented as mean ± SEM. Statistical analyses were performed using (B) a multiple t-test and (C and D) an unpaired student's t-test. APC = allophycocyanin, ePPE = external porcine pancreatic elastase, GP = glycoprotein, HCT = Haematocrit, HGB = Haemoglobin, IL4R = interleukin 4 receptor, MFI = mean fluorescence

### GPIIbα in abdominal aortic aneurysm formation

intensity, MPV = mean platelet volume, PCT = plateletcrit, PDW = platelet distribution width, PE = phycoerythrin, plts = platelets, RBCs = red blood cells, WBCs = white blood cells, WT = wildtype.

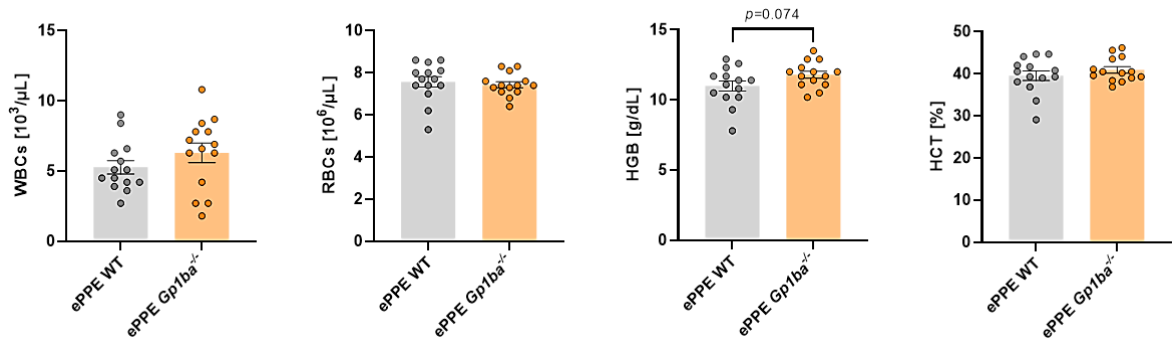

**Figure S2. GPIIbα deficiency in mice does not affect white or red blood cells counts following ePPE surgery.** WBC and RBC counts, respectively HGB and HCT of ePPE operated WT and *Gp1ba*<sup>-/-</sup> ($n = 14$) mice at day 28 post surgery. Data are presented as mean ± SEM. Statistical analyses were performed using an unpaired student's t-test. ePPE = external porcine pancreatic elastase, HCT = Haematocrit, HGB = Haemoglobin, RBCs = red blood cells, WBCs = white blood cells, WT = wildtype.

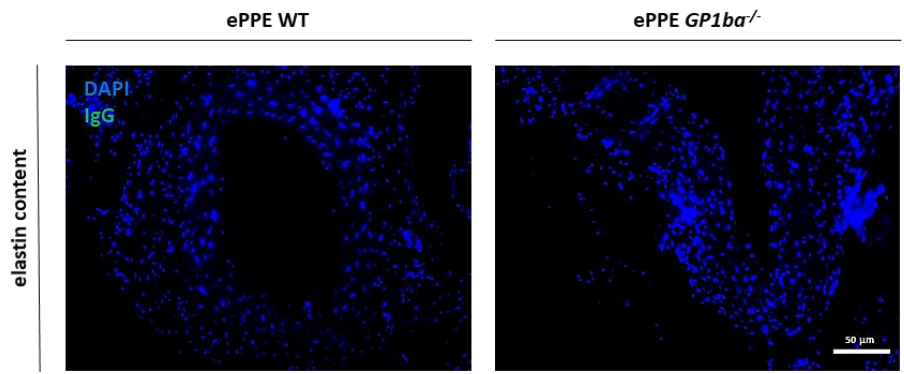

**Figure S3. IgG controls of immunofluorescence images of murine aortic tissue after ePPE induced AAA formation.** Representative images of IgG controls from elastin immunofluorescence (IgG/Cy3; green) staining of murine aortic tissue of ePPE-operated WT and *Gp1ba*<sup>-/-</sup> ($n = 5$) mice at day 28 post surgery. Nuclei were stained with DAPI (blue). ePPE = external porcine pancreatic elastase, WT = wildtype.

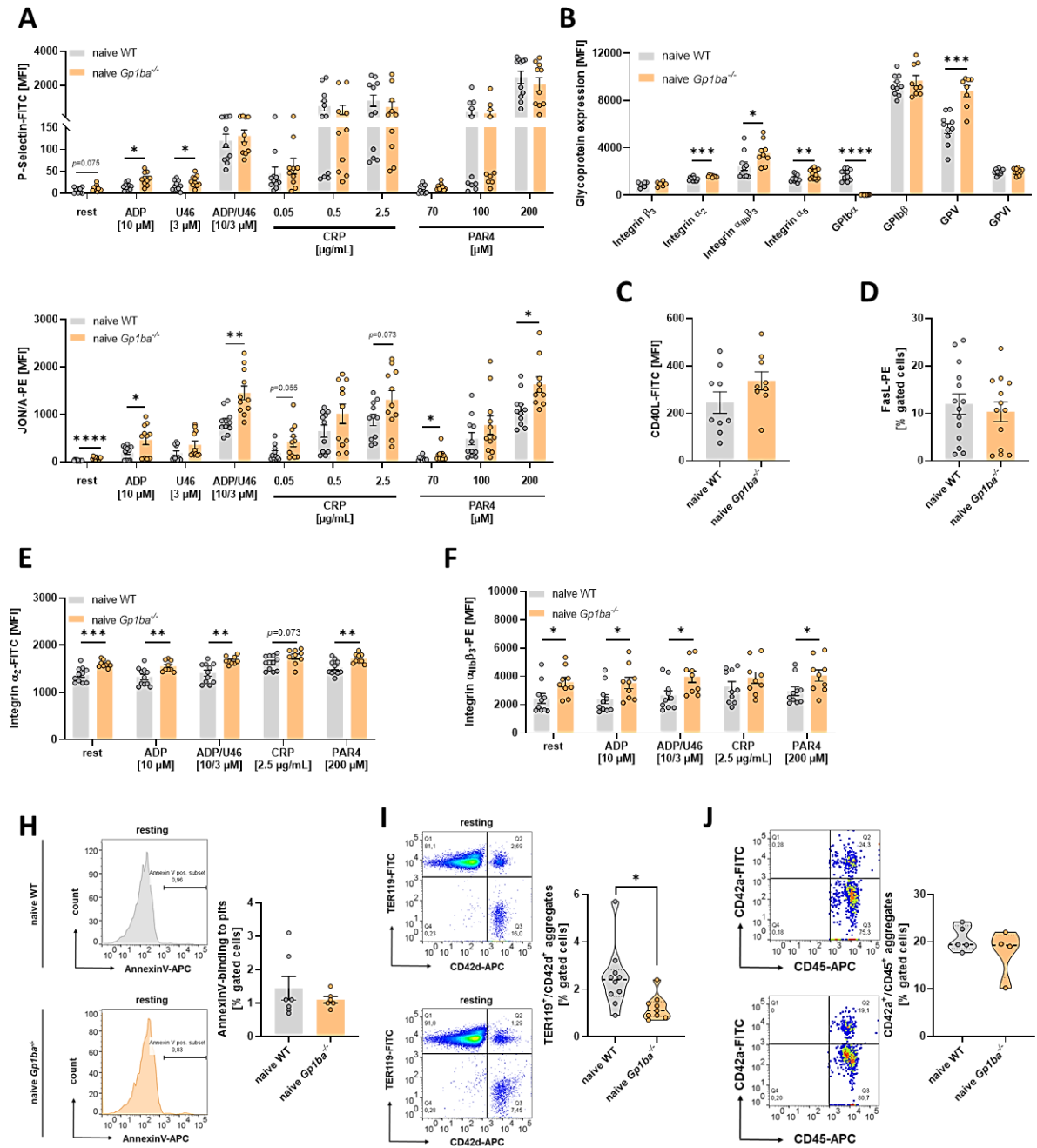

**Figure S4. Loss of GPIIb is associated with elevated glycoprotein externalisation as well as a sustained platelet hyperactivity.** (A–J) Washed whole blood of naïve WT and *Gp1ba*<sup>-/-</sup> mice was analysed via flow cytometry. (A) Platelet degranulation (P-selectin-FITC) and integrin $\alpha_{IIb}\beta_3$ activation (JON/A-PE) ($n = 10–11$) were determined. (B) Platelet surface exposure of integrin $\beta_3$, integrin $\alpha_2$, integrin $\alpha_{IIb}\beta_3$, integrin $\alpha_5$, GPIIb, GPIbβ, GPV and GPIV were analysed ($n = 6–16$). Platelets were stimulated with indicated agonists. (C and D) Surface exposure of (C) CD40L ($n = 9$) and (D) FasL ($n = 13–14$) on the platelet surface under resting conditions. (E and F) Externalisation of (E) integrin $\alpha_2$ ($n = 9–11$) and (F) integrin $\alpha_{IIb}\beta_3$ ($n = 9–11$) on the platelet surface. Platelets were stimulated with indicated agonists. (H) PS exposure of platelets was analysed by Annexin V binding ($n = 6–7$). (I and J) Aggregate formation of platelets with (I) RBCs ($n = 9–10$) and (J) leukocytes ($n = 4–5$) was analysed as double-positive events for CD42d (platelet maker, GPV) and TER119 (RBC marker), respectively CD45 (leukocyte marker). Data are presented as mean ± SEM. Statistical analyses

were performed using (**A**, **B**, **E** and **F**) a multiple t-test and (**C**, **D**, **H**, **I** and **J**) an unpaired student's t-test. ADP = Adenosine diphosphate, APC =allophycocyanin, t. CRP = collagen-related peptide, FITC = fluorescein isothiocyanate, GP = glycoprotein, MFI = mean fluorescence intensity, PAR4 = protease-activated receptor 4 activating peptide, PE = phycoerythrin, U46 = thromboxane A2 analogue U46619, WT = wildtype.

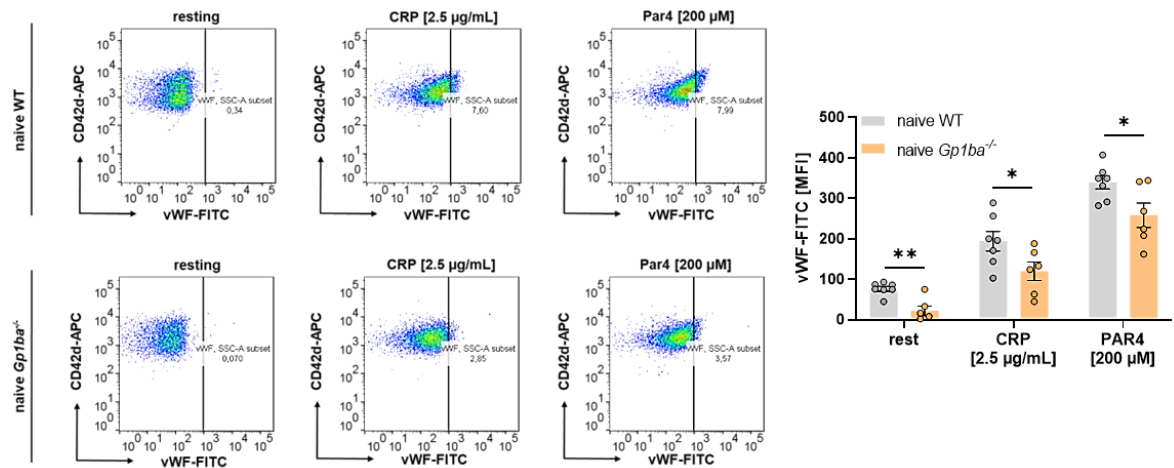

**Figure S5. Genetic replacement of the GPIIbα receptor results in reduced VWF binding to platelets.** Washed whole blood of naive WT and *Gp1ba*<sup>-/-</sup> mice was analysed for VWF binding to platelets ($n = 6-7$) using flow cytometry. Platelets were stimulated with indicated agonists. Data are presented as mean ± SEM. Statistical analyses were performed using (B and C) a multiple t-test. FITC = fluorescein isothiocyanate, ePPE = external porcine pancreatic elastase, MFI = mean fluorescence intensity, VWF = Von Willebrand factor, WT = wildtype.
