## Supplementary material for "Dysregulated Platelet GPIbα–VWF Signalling in Abdominal Aortic Aneurysm formation and Progression": Table 1

**Table S1. Clinical parameter of AAA patients.** Continuous data are represented as mean ± SEM. Dichotomous data are represented as absolute frequency with percentage (%). Control whole blood samples were collected from anonymized healthy donors ($n = 11$, average age: $65.5 \pm 1.3$ years) from the blood bank of the University Hospital Duesseldorf.

AAA = abdominal aortic aneurysm; aHT = arterial hypertension; CHD = coronary heart disease; SAPT = single anti-platelet therapy; T2D: type 2 diabetes.

|  | Age<br>[years] | Male sex | History of<br>smoking | CHD | aHT | T2D | Obesity | Statins | SAPT | Antihypertensive<br>drugs |
| --- | --- | --- | --- | --- | --- | --- | --- | --- | --- | --- |
| **AAA** | 75.9 | 81.2% | 37.5% | 25.0% | 75.0% | 18.75% | 0% | 100% | 75.0% | 87.5% |
| **(n =16)** | $\pm 2.1$ | (13) | (6) | (4) | (12) | (3) | (0) | (16) | (12) | (14) |
